# The use of thermostable fluorescent proteins for live imaging in *Sulfolobus acidocaldarius*

**DOI:** 10.1101/2024.06.16.599207

**Authors:** Alejandra Recalde, Jasmin Abdul Nabi, Pierre Junker, Chris van der Does, Jana Elsässer, Marleen van Wolferen, Sonja-Verena Albers

## Abstract

Among hyperthermophilic organisms, *in vivo* protein localization is challenging due to the high growth temperatures that can disrupt proper folding and function of mostly mesophilic-derived fluorescent proteins. While protein localization in the thermophilic model archaeon *S. acidocaldarius* has been achieved using antibodies with fluorescent probes in fixed cells, the use of thermostable fluorescent proteins in thermophilic archaea has so far been unsuccessful.

Given the significance of live protein localization in the field of archaeal cell biology, we aimed to identify fluorescent proteins for use in *S. acidocaldarius*. To achieve this, we expressed various previously published and optimized thermostable fluorescent proteins along with fusion proteins of interest and analyzed the cells using flow cytometry and (thermo-) fluorescent microscopy. Of the tested proteins, Thermal Green Protein (TGP) exhibited the brightest fluorescence when expressed in Sulfolobus cells. By optimizing the linker between TGP and a protein of interest, we could additionally successfully fuse proteins with minimal loss of fluorescence. TGP-CdvB and TGP-PCNA1 fusions displayed localization patterns consistent with previous immunolocalization experiments. These initial results in protein localization in *S. acidocaldarius* at high temperatures, combined with recent advancements in thermomicroscopy, open new avenues in the field of archaeal cell biology. This progress finally enables localization experiments in thermophilic archaea, which have so far been limited to mesophilic organisms.

## Introduction

Since its discovery in 1962, the Green Fluorescent Protein (GFP) from *Aequorea victoria* and its derivatives have proven to be invaluable tools for observing microscopic life (Pakhomov & Martynov, 2008; Phillips, 2001; Shimomura et al., 1962). When fused to another protein, fluorescent proteins (FPs) allow the intracellular localization of these and the visualization of processes in living cells, such as cell division, translation, and DNA replication. In archaea, the use of FPs has proven to be challenging because of the extreme growth conditions that many of the model organisms live in.

To date, their use in archaea has been limited to mesophilic organisms such as *Haloferax volcanii*. In this organism, specific point mutations in GFP resulted in a protein that is stable under high salinity conditions (Reuter & Maupin-Furlow, 2004) and has successfully been used to study various cellular processes, including cell division, S-layer synthesis, DNA replication, cell motility, and cell shape (Bisson-Filho et al., 2018; Ithurbide et al., 2022, 2024). Additionally, the creation of an autofluorescence-free *H. volcanii* strain enabled the establishment of live single-molecule microscopy (Turkowyd et al., 2020).

The use of FPs in methanogens has been restricted by the anoxic conditions these organisms require, as oxygen is necessary for the proper maturation of the fluorophore (Heim et al., 1994). However, the development of oxygen-independent fluorescence-activating and absorption-shifting tag (FAST) has proven useful (Hernandez & Costa, 2022). FAST tags exhibit fluorescence upon binding to specific fluorogens. Various FAST variants, with affinity to different ligands, are commercially available (Tebo et al., 2021).

For thermophiles, such as members of the Sulfolobales, the lack of FPs that function at high temperatures has necessitated the use of immunolocalization on fixed cells to visualize cell division proteins (Cezanne et al., 2023; Lindås et al., 2008; Samson et al., 2008; Tarrason Risa et al., 2020). In studies on the thermophilic bacterium *Thermus thermophiles*, superfolder GFP (sfGFP), a variant of GFP was used at 70°C to localize GroES in the cell (Cava et al., 2008). sfGFP was generated to have a high chemical stability and to fold efficiently even in the presence of misfolded fusion proteins (Pédelacq et al., 2006). So far, it was not expressed in any thermophilic Archaea, in this study we therefore tested several presumed thermostable FPs in *S. acidocaldarius*.

The thermostable FP eCGP123 was the first FP that was successfully expressed in *Sulfolobus acidocaldarius*, allowing the visualization of biofilms (Henche et al., 2012). It was developed through direct evolution of a Consensus Green Protein (CGP) (Kiss et al., 2009). Further engineering resulted in the so-called Thermal Green Protein (TGP), which is more stable and less prone to aggregation (Close et al., 2015). TGP retained fluorescence *in vitro* after prolonged incubation at 90°C, but it has not been tested in thermophilic organisms *in vivo* (Close et al., 2015). Recently, point mutations in TGP led to the development of an enhanced version (TGP-E) (Anderson et al., 2023).

FPs with different colors enable the simultaneous labeling of various structures within a cell. In addition to green FPs, we therefore tested a few thermo-optimized yellow FPs. Mutagenesis of sfGFP and TGP, respectively, led to the development of Superfolder YFP (sfYFP) (Pédelacq et al., 2006), as well as the Yellow Thermo Protein (YTP) and its enhanced version (YTP-E) (Anderson et al., 2023). Moreover, the yellow hyperfolder YFP (hfYFP), derived from mGreenLantern, and its monomeric version mfYFP are highly thermostable, resistant to acidic conditions and to chemical agents used to fixed cells, such as PFA and OsO_4_ (Campbell et al., 2022).

In recent years, there has been growing interest in the cell biology of archaea. Among halophiles, significant developments in cell biology, including the use of diverse fluorescent proteins, have facilitated live microscopy of intracellular structures, comparable to that in bacteria (Bisson-Filho et al., 2018; Ithurbide et al., 2024; van Wolferen et al., 2022). Additionally, the optimization of thermomicroscopy has allowed the visualization of thermophilic organisms, such as the model organism *S. acidocaldarius*, under their natural growth temperatures (Charles-Orszag et al., 2021; Mora et al., 2014; Pulschen et al., 2020). Nevertheless, especially amongst thermophiles the field of cell biology is still lagging behind, mainly due to the absence of functional thermostable FPs. In this study, we therefore explore the use of TGP and other green and yellow thermostable FPs in *S. acidocaldarius*. Fusion proteins of TGP with CdvB and PCNA1 confirmed their previously published localization. In addition, in-gel fluorescence of the proteins enabled straightforward detection without the requirement for Western blotting.

## Results

### Finding a suitable fluorescent protein for use at high temperatures

In our search for thermostable FPs suitable for localization experiments in *S. acidocaldarius*, we first tested three previously published thermostable green FPs: sfGFP (Pédelacq et al., 2006), eCGP123 (Kiss et al., 2009) and TGP (Close et al., 2015) (Table 1). Codon optimized genes were first expressed using our previously published FX expression plasmids (van der Kolk et al., 2020) by inducing the cells with xylose. To measure their brightness, we performed flow cytometry on the expression cultures (Fig. 1AB, Fig. S1AB). Induction with either 0.2% or 1% xylose gave similar results. Additionally, both preheated samples (kept at around 75°C until injection) and cells transported at room temperature did not show any significant difference in fluorescence (Fig. S1A-B). Cells expressing either sfGFP or eCGP123 showed only slightly increased fluorescence compared to the autofluorescence of cells expressing an empty plasmid (Fig1A, FigS1A). TGP exhibited the brightest fluorescence when expressed in *S. acidocaldarius*.

**Figure 1.**
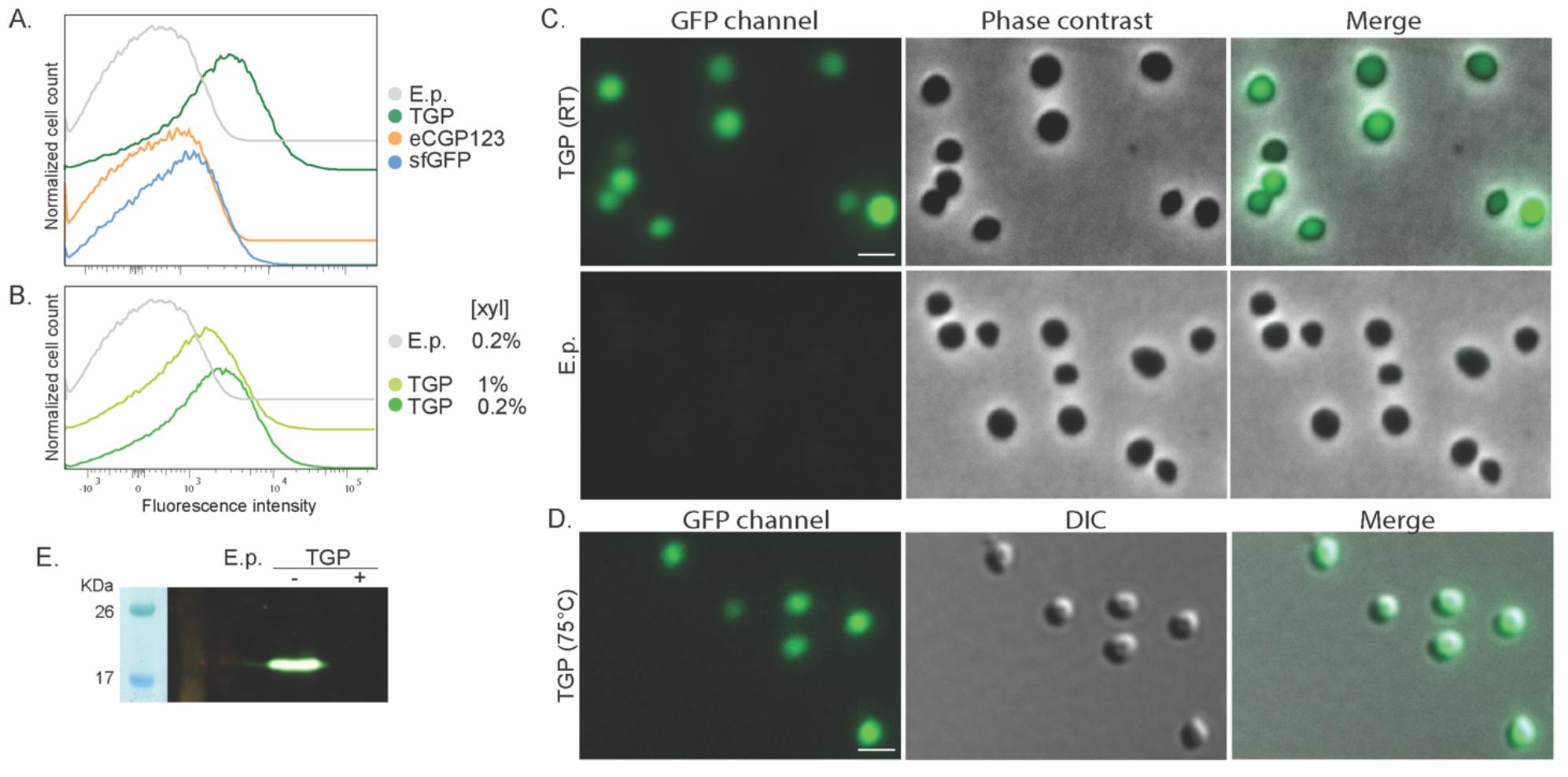
Flow cytometry on *S. acidocaldarius* cells expressing (A) different green fluorescent proteins induced with 0.2% xylose and (B) TGP induced with different xylose concentrations. Total number of events per sample: 100.000. (C) Fluorescent microscopy, phase contrast and merged images of cells at RT (D) Fluorescent microscopy, DIC and merged images of cells at 75°C. Scale bar: 2 μm. (E) In-gel fluorescence of TGP, when samples were boiled (+) or not (-) in SDS containing buffer. E.p.: Empty plasmid, RT: Room temperature.

In line with our flow cytometry experiments, cells expressing TGP were clearly more fluorescent than cells expressing either sfGFP or eCGP123 when visualized at RT using fluorescence microscopy (Fig. 1C and S1C). Importantly, similarly bright fluorescence could be observed when visualized at 75°C using the VAHEAT system, making TGP well-suited for live imaging in *S. acidocaldarius* (Fig 1D).

TGP could also be visualized in-gel using our fluorescence imager. For this, whole cell lysates were loaded on an SDS-gel. Samples that were boiled for 10 min did not show any fluorescence, whereas unboiled samples revealed a nicely fluorescent band at the expected height of TGP (Figure 1E).

We have thereby successfully used TGP as a fluorescent marker in *S. acidocaldarius*, facilitating flow cytometry, fluorescence microscopy and in-gel fluorescence. Notably, its fluorescence remains similarly bright under the optimal growth conditions of *S. acidocaldarius*, enabling dynamic live imaging.

### Position of the companion protein and use of linkers affected fluorescence of TGP

Because our ultimate goal would be to efficiently localize proteins in *S. acidocaldarius*, we fused TGP both N- and C-terminally to the cytosolic protein LacS and an HA tag, linked with a short linker (GGGSGGG, Table 2). Flow cytometry revealed that all fusions resulted in decreased fluorescence when compared to TGP alone, particularly noticeable when TGP was fused at the C-terminus of the protein (Fig. 2A). The latter could also be observed in fluorescence microscopy (Fig. 3D). Because linkers can significantly affect protein activity or even increase expression yields (Chen et al., 2013; Ithurbide et al., 2024; Waldo et al., 1999), we decided to test four different previously published linkers: a linker designed for rapid protein folding assays with fluorescent proteins, which we renamed as thermolinker (Waldo et al., 1999); along with a flexible-, semi-rigid- and rigid linker (Chen et al., 2013) (Fig. 2B). Their sequences are summarized in Table 2.

**Figure 2.**
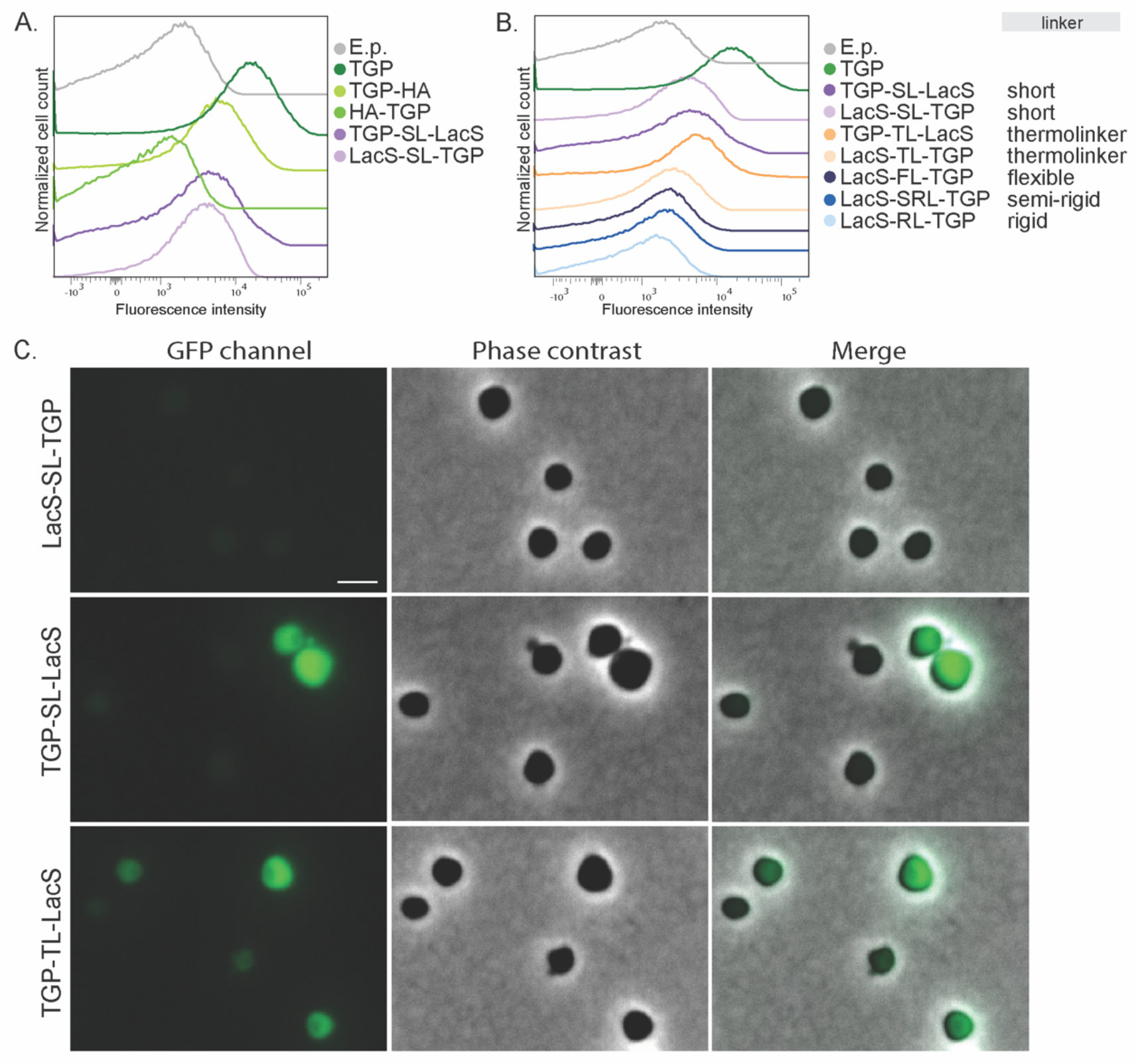
(A) Fluorescence intensity of *S. acidocaldarius* cells expressing different TGP-LacS fusions at RT and 0.2% xylose. (B) Influence of different linkers in the intensity of fluorescence of TGP-LacS fusions. Number of events for FC: 100.000. (C) Fluorescent microscopy, phase contrast, and merged images of cells expressing different TGP-LacS fusions. Adjustment of the histogram curve was done according to the microscopy image of TGP in Figure 1C. Scale bar: 2 μm. E.p.: Empty plasmid.

**Figure 3.**
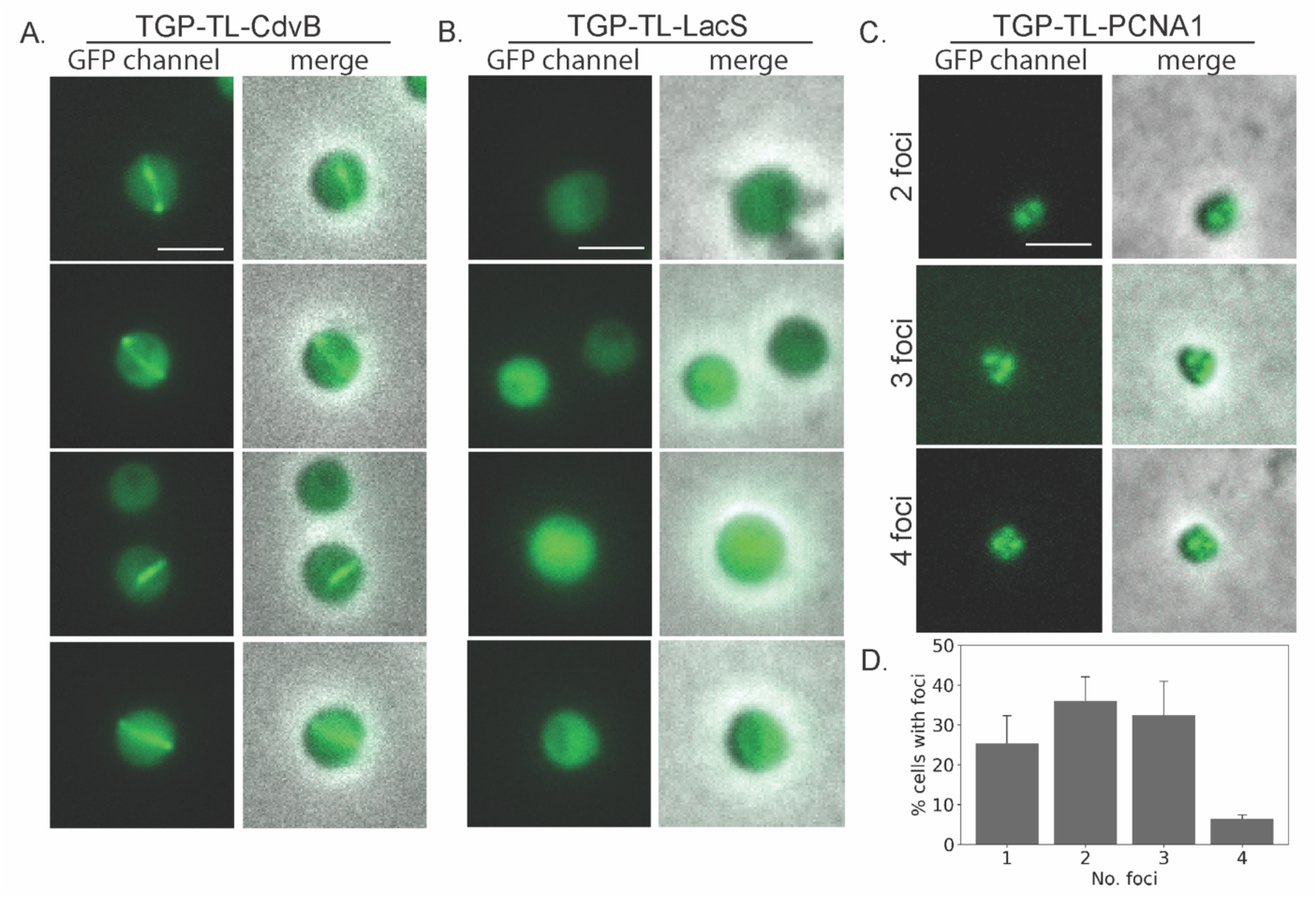
Localization of different TGP fusion proteins in *S. acidocaldarius*. Fluorescence microscopy images and a merge with phase contrast images of (A) CdvB rings (B) LacS and (C) PCNA. (D) Number of foci per cell in PCNA localization (three pictures were used for counting foci, n=48). Scale bar: 2 μm.

Fusion proteins linked with the thermolinker (TL), semi-rigid linker (SRL) and the flexible linker (FL) exhibited slightly higher fluorescence compared to the fusion with the original short linker (Fig. 2C). Especially TGP-TL-LacS showed an increased fluorescence as was also evident in fluorescence microscopy (Fig.2C and 2D).

We attempted to enhance brightness and stability of TGP-fusion proteins by introducing point mutations to alter the pKa. Additionally, a cysteine-free version of the TGP was created, considering the rarity of disulfide bonds in archaea. Unfortunately, none of these mutations yielded fluorescence comparable to the original TGP (Fig. S2). Therefore, we continued using the original TGP in combination with a thermolinker, preferably at the C-terminus of the protein.

To facilitate cloning in future studies, we created plasmid pSVAara_TGP-TL_FXStop (Fig. S3, Table S1), based on our previously published set of plasmids (van der Kolk et al., 2020). It harbors the arabinose promoter used in this study, which is responsive to arabinose and xylose (van der Kolk et al., 2020). The plasmid contains *tgp* and the thermolinker. Moreover, *lacI/Z* genes flanked by *Nco*I and *Xho*I allow easy cloning of a gene of interest using blue-white screening. Alternatively, SapI restriction sites would allow for FX-cloning (Geertsma & Dutzler, 2011; van der Kolk et al., 2020).

### Imaging cell division and replication foci *in vivo*

Previously, cell division proteins of *Sulfolobus* species have been rewarding targets for localization, providing a clear localization pattern at mid-cell. Especially ESCRT-III homologs CdvB, CdvB1 and CdvB2 form distinct rings. However, due to the absence of thermostable FPs, immunolocalization has been the only method to visualize these proteins, involving fixing and permeabilizing cells (Lindås et al., 2008; Pulschen et al., 2020; Samson et al., 2008; Tarrason Risa et al., 2020).

To test if we could observe similar localization patterns using TGP, we expressed a TGP-TL-CdvB fusion in *S. acidocaldarius*. It was previously shown that inhibiting the proteasome in *S. acidocaldarius* impairs the degradation of CdvB rings (Tarrason Risa et al., 2020), allowing them to be visible for a longer duration. Therefore, we synchronized the cells with acetic acid and subsequently added the proteasome inhibitor bortezomib. As previously observed with immunolocalization, CdvB rings could be observed at midcell after bortezomib arrest (Figure 3A). Control cells expressing TGP-TL-LacS showed diffused localization (Figure 3B).

As the second target with a distinct localization pattern, we chose the DNA sliding clamp protein PCNA. In Sulfolobales, three PCNA subunits form a heterotrimeric structure (Acharya et al., 2021; Williams et al., 2006). Immunolocalization of PCNA1 has allowed replisome positioning in *S. acidocaldarius*, revealing one to four foci in the cells during G1 (Gristwood et al., 2012). To test if TGP could similarly be used for PCNA and replisome localization, we expressed TGP-TL-PCNA1 (Saci_0826) under the control of its native promotor. After synchronization, cells were observed under the microscope. Consistent with previous findings (Gristwood et al., 2012), we observed cells with a variable number of foci (between 1 and 4) (Figure 3C and D).

We have thereby, for the first time, expressed functional fluorescent fusion proteins *S. acidocaldarius*, enabling the localization of proteins in living cells.

### Other fluorescent proteins

Ideally, we would like to have a diverse color palette of FPs for *S. acidocaldarius*, similar to those available for Haloferax (Ithurbide et al., 2024) and Bacteria. To achieve this, we tested a set of previously published thermostable yellow FPs (Table 1). These included: sfYFP, hfYFP, mfYFP, YTP and YTP-E.

Similar to sfGFP, sfYFP did not show any fluorescence when expressed in *S. acidocaldarius* cells (Figure S4). YTP and YTP-e were promising candidates, given the fact that they are derived from TGP (Anderson et al., 2023). Unexpectedly, both YTP and YTP-E exhibited low fluorescence under our tested conditions (Figure S4).

Two other yellow proteins, hfYFP and its monomeric variant mfYFP, were reported as good alternatives to enhanced YFP (e-YFP) because of their higher thermostability and the fact that their excitation spectra do not overlap with that of enhanced GFP (e-GFP) (Campbell et al., 2022). Both hfYFP and mfYFP were less fluorescent than TGP under our tested conditions. However, fluorescence could still be observed when expressing monomeric mfYFP (Figure 4A). Similar to cells expressing TGP, in-gel fluorescence was observed when loading whole cells expressing hfYFP or mfYFP on a gel, though with less intensity (Figure 4B). We also tested the use of commercially available GFP antibodies and were able to detect both hfYFP and mfYFP, but not TGP (Fig. 4C). This provides an additional method to detect these proteins without the need for a tag.

**Figure 4.**
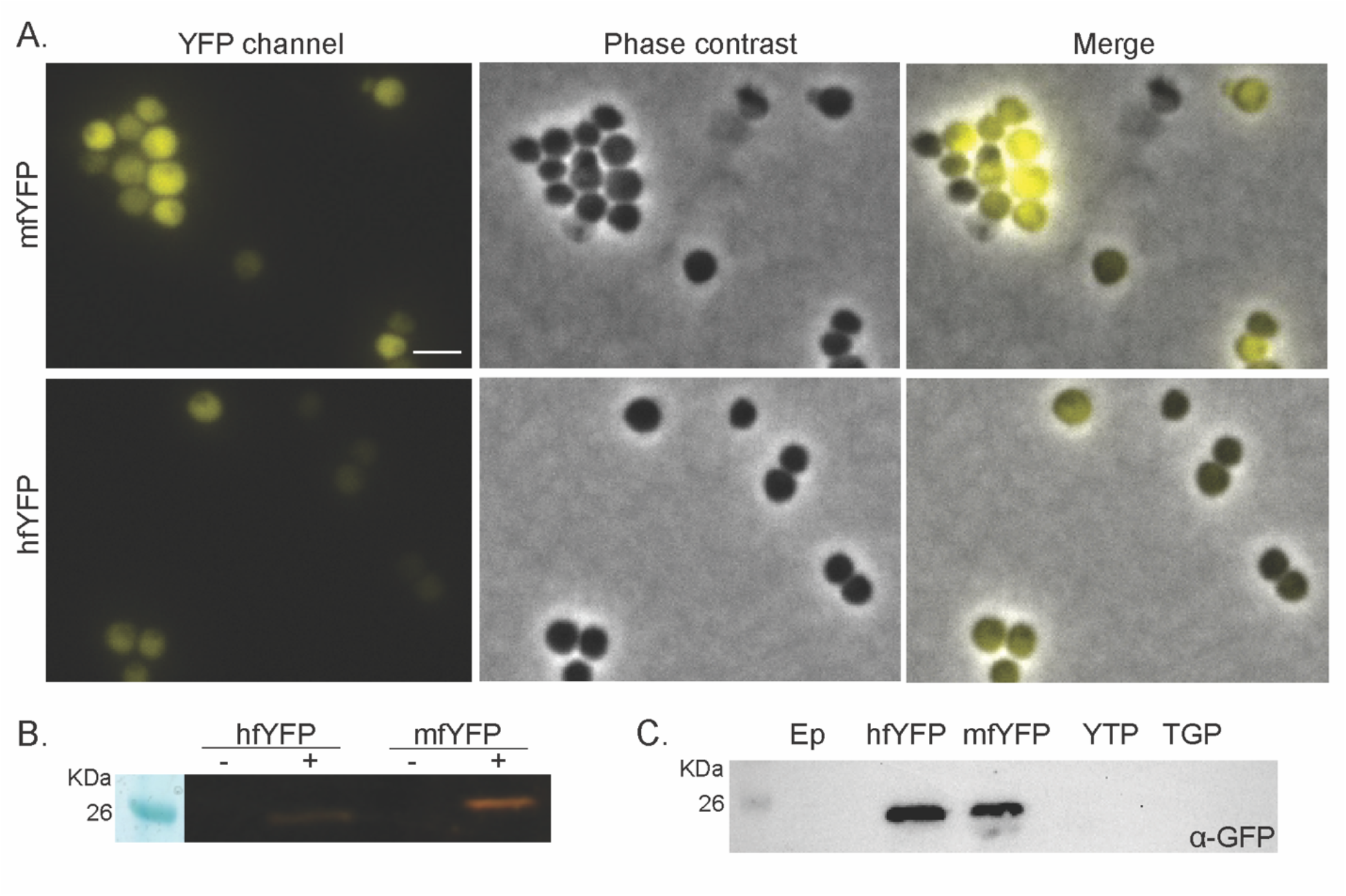
Expression of yellow FPs in *S. acidocaldarius*. (A) Fluorescent microscopy, phase contrast, and merged images of *S. acidocaldarius* cells at RT expressing mfYFP and hfYFP, respectively. Scale bar: 2 μm. (B) In gel fluorescence of hfYFP and mfYFP, samples were boiled (+) or not (-) in SDS containing buffer. (C) Western blot with anti-GFP antibodies.

With mfYFP, we thereby now have an additional thermostable FP with a different excitation wavelength than TGP. Optimization of this protein and the search for thermostable FPs in other colors will expand our toolbox for live fluorescent thermomicroscopy in *S. acidocaldarius*.

## Discussion and Conclusion

Recent developments in thermomicroscopy significantly contributed to the field of live microscopy of thermophilic archaea, including *S. acidocaldarius*. These advancements have enabled the visualization of dynamic cellular processes in real-time, such as motility and cell division (Charles-Orszag et al., 2021, 2023; Pulschen et al., 2020). With the availability of easy-to-use thermomicroscopy, the development of thermostable fluorescent markers and proteins has become increasingly desirable. While it is possible to utilize dyes for staining components such as DNA or the membrane at high temperatures (Cezanne et al., 2023), the visualization of proteins in Sulfolobales has so far relied on immunostaining (Bisson-Filho et al., 2018; van Wolferen et al., 2022).

In this study, we tested different thermostable FPs, linkers, and fusions to identify FPs for studying the cell biology of living *S. acidocaldarius* cells. Our findings highlight the successful use of TGP and mfYFP in *S. acidocaldarius* for live imaging. Furthermore, in-gel fluorescence, even under denaturing conditions, allowed easy protein detection on SDS-gels, eliminating the need for western blotting and specific antibodies. When fusing TGP to CdvB and PCNA1, the fusion proteins revealed distinct localization patterns that were previously observed in immunolocalization studies (Gristwood et al., 2012; Samson et al., 2008), demonstrating the effectiveness of TGP in visualizing intracellular proteins. Surprisingly, TGP-based YTP and E-YTP did not exhibit fluorescence despite their thermostability. Possibly due to the fact that, although these proteins are more thermostable than the TGP, they have been reported to possess lower quantum yields (Anderson et al., 2023).

So far, we could only successfully localize proteins that were N-terminally tagged with TGP. Additionally, mfYFP showed significantly less bright fluorescence compared to TGP. Future optimization strategies, such as the use of error-prone PCR libraries, could further enhance the brightness and stability of TGP and mfYFP in *S. acidocaldarius*, as it was done in thermophilic bacteria (Frenzel et al., 2018). In addition, the ongoing search for FPs in alternative colors will, in the future, hopefully, enable the colocalization of diverse proteins within the same cell, thereby enhancing our understanding of more complex cellular dynamics and interactions. We anticipate that the utility of TGP and mfYFP can extend beyond *S. acidocaldarius* to other thermophilic archaea, providing valuable insights into protein localization and dynamics within several thermophilic organisms.

## Material and methods

### Strains, media, transformation and growth conditions

*Sulfolobus acidocaldarius* MW001 was grown in Brock medium (Brock et al., 1972) supplemented with 0.1% (*w/v*) N-Z amine (Sigma-Aldrich^®^, Merk KGaA, Burlington, MA, USA), 0.2% (*w/v*) sucrose, and 0.01 mg/mL uracil in the case of uracil autotrophs. Cells were grown in shaking conditions at 120 rpm and 75°C. For expression cultures, 0.2 or 1% of xylose was added in the exponential phase (OD600 0.3-0.5). Subsequently, the cultures were incubated at 75°C for 4 hours or overnight. For plates, two times concentrated Brock medium supplemented with 6 mM CaCl_2_, 20 mM MgCl_2_, 0.2% N-Z-amine (*w/v*), and 0.4% dextrin (*w/v*) was pre-warmed and mixed in the same volume of freshly boiled 1.4% generate (Carl Roth, Karlsruhe, Germany).

Competent cells were prepared as previously described (Wagner et al., 2012; Ye et al., 2022). For transformation, 200 ng of methylated plasmid DNA was mixed with competent cells. Electroporation was done in a 1 mm cuvette using a Gene Pulser Xcell (BioRad, München, Germany) with a constant time protocol with input parameters 1.5 kV, 25 μF and 600 Ω. 400 μL of recovery solution (Basic Brock medium without pH adjustment) was added and cells were recovered for 30 min shaking at 300 rpm and 75°C. Afterward, 100 μL of cell suspension was plated on plates without uracil and incubated inside a humidity chamber for 6-7 days.

Competent *Escherichia coli* DH5α, Sure2 or ER1821 (NEB) used for cloning and methylation of plasmid DNA respectively, were grown in at 37°C LB supplemented with the appropriate antibiotics.

### Plasmids construction

All plasmids and primers are listed in Table S1 and S2.

Different codon optimized *tgp* variants (low and high pKa and a cysteine free version), codon optimized *hf-yfp* as well as linker sequences were ordered from GenScript. Genes for FPs and linkers were fused using overlap PCR. Resulting fragments were cloned into the listed expression plasmids using *Nco*I and *Apa*I. Promoter sequences were cloned using *Sac*II and *Nco*I. The TGP derived variants YTP (TGP H193Y) and YTP-e (Q65E, H193Y), as well as the monomeric version of hfYFP, mfYFP (hfYFP S147P, L195M, V206K) were created by subsequent side directed point mutagenesis.

If not stated otherwise, genes of interest (*lacS, cdvB* and *pcna1)* were amplified using the primers listed in Table S2 and cloned into the expression plasmids (van der Kolk et al., 2020) with *Nco*I and *Xho*I (New England Biolabs GmbH, Frankfurt am Main, Germany). Correct plasmids were methylated by transforming them to ER1821 containing pM.EsaBC4I, expressing a methylase (NEB).

### Flow cytometry

Samples from expression cultures were taken and kept warm and away from light exposure in a container until injection into a Becton-Dickinson Fortessa flow cytometer equipped with a green laser (488 nm). Filters used for detection were FL12 – 450/50 and FL13 – 530/30 – 550 LP. The number of events recorded per sample was 100,000. Analysis was carried out using FlowJo™ software v10.10.

### Cell synchronization

Cells from expression cultures (OD_600_ of 0.1-0.2) were synchronized using acetic acid as described previously (Tarrason Risa et al., 2020). Expression was induced from the beginning of synchronization with 0.1% xylose. Brock medium for washing steps and final resuspension always included xylose. Samples were taken 80-100 min after washing away the acetic acid. For proteasome arrest, 80 min after release, 10 mM bortezomib was added and cells were grown for 30 min at 75°C. Cells were then washed twice with Brock media, placing them back in the incubator for 10 min in between, and incubated again at 75°C. Samples were taken after 2, 5, 7 and 10 min.

### Fluorescent microscopy at room temperature and 75°C

For imaging, 3 μL of cell suspension was spotted on an agarose pad (1% agarose in basic Brock medium), air dried, and imaged with an Axio Observer Z1; Zeiss microscope equipped with a Plan-Apochromat 100x 1.40 Oil Ph3 M27 objective. At least three fields were imaged per sample and experiments were repeated three times. Exposure time was of 1 s for GFP channel (450-490/500-550) and 500 ms in YFP channel (490-510/520-550).

For thermomicrosocpy, 1 mL of culture was taken and added to a VaHeat substrate chamber, and closed with a small glass slide (Interherence GmbH, Germany). The sample was then heated to 75°C for 5 minutes before imaging. Images were taken with an Axio Observer Z1; Zeiss microscope equipped with a Plan-Apochromat 100x 1.40 Oil DIC M27 objective. The exposure time was 1 s for the GFP channel (450-490/500-550).

### SDS-page, western blot and in gel fluorescence

Pellets from 10 mL cell cultures were resuspended in 1x PBS to a theoretical OD of 10. The samples were mixed with SDS-dye (50 mM Tris-HCl pH 6.8, 2% glycerol, 2% DTT, 0.0004% Bromophenol blue, 2% SDS) and boiled, or not, for 10 minutes at 99°C. Cell lysates were then run on 15% sodium dodecyl sulfate-polyacrylamide gel via electrophoresis. Gels were either stained with Coomassie (25% v/v isopropanol, 10% acetic acid, 0.05% Coomassie R) or blotted onto a polyvinylidene fluoride (PVDF) membrane using the semi-wet Western blot Trans-Blot Turbo Transfer System (BioRad). Blocking was done overnight at 4°C using a 0.1% I-Block (Thermo Fisher Scientific) solution in PBS-T (PBS + 0.1% Tween 20). The primary antibody α-HA (rabbit) (Sigma-Aldrich) was used in a 1:10,000 dilution and incubated for 4 hours at 4°C. Primary antibody α-GFP raised in rabbit (Sigma) was used in a 1:10,000 dilution and incubated overnight at 4°C. The membrane was washed 3 times with PBS. The secondary antibody, anti-rabbit HRP coupled (1:10,000 dilution, Sigma-Aldrich), was then applied and incubated for 3 h. Chemiluminescence reaction was initiated by adding HRP substrate (Clarity Max Western ECL Substrate, BioRad). Signals for the Western blot and in-gel fluorescence were acquired using the iBright FL1500 imaging system (Invitrogen). For in-gel fluorescence, the gel was imaged immediately after protein separation.

## Authors contributions

AR and S-VA conceived the study. JAN and PJ constructed the plasmids and performed imaging acquisition under the supervision of AR. AR performed cloning, cell synchronization, fluorescent microscopy, flow cytometry, and SDS gel and western blot experiments. JE created early plasmids and performed preliminary fluorescence microscopy under the supervision of MvW. CvdD designed the TGP variants. AR and MvW wrote the manuscript.

## Acknowledgment

We acknowledge support by the Open Access Publication Fund of the University of Freiburg.

