## Supplementary Material for "The use of thermostable fluorescent proteins for live imaging in *Sulfolobus acidocaldarius*"

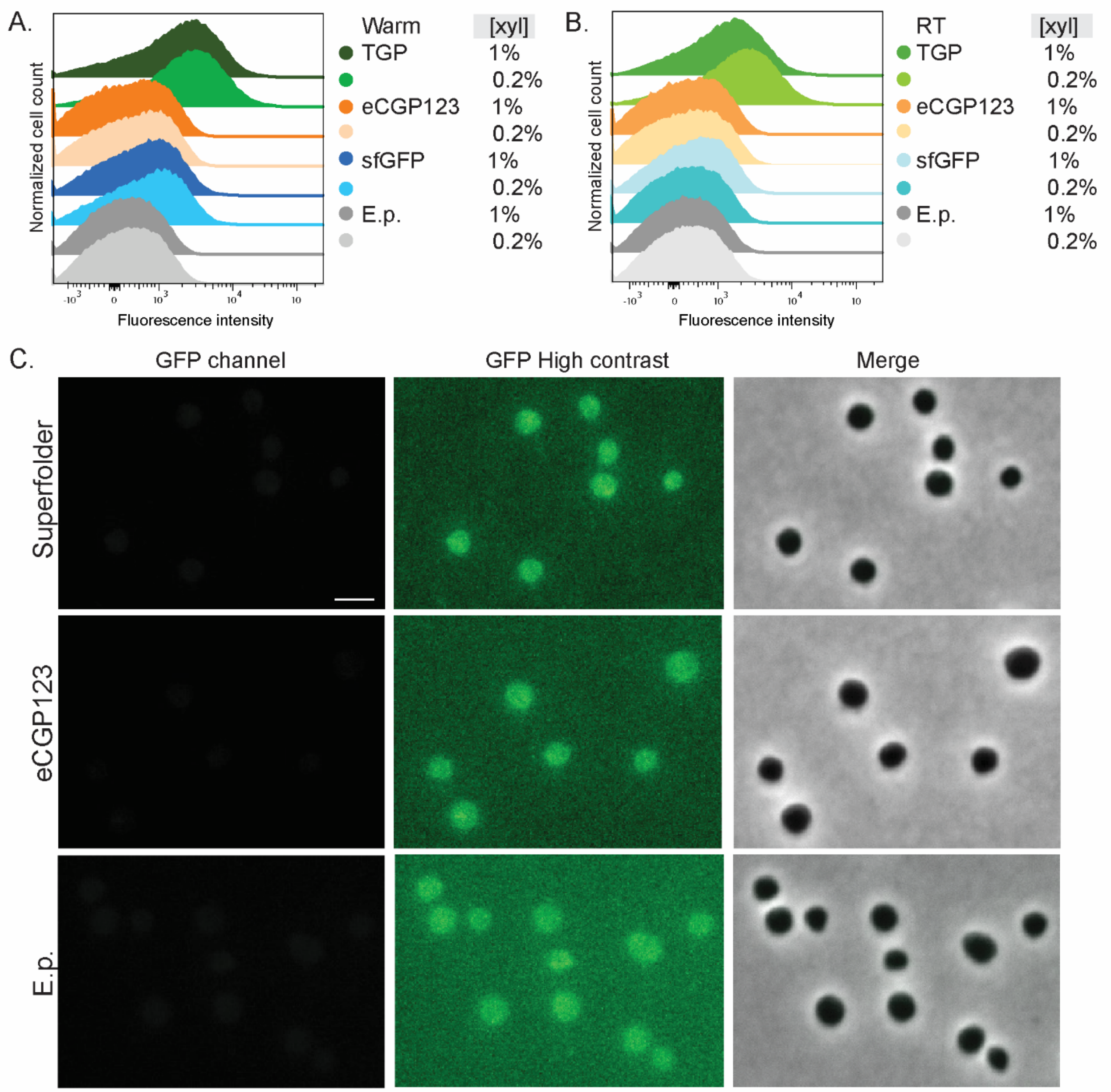

Figure S1. *S. acidocaldarius* expressing different green FPs. Fluorescence intensity in flow cytometry of preheated samples (warm) (A) and samples transported at room temperature (B) induced with 0.2 or 1% xylose. Number of events for FC: 100.000. (C) Fluorescent microscopy, phase contrast and merge with phase contrast images of cells at 0.2% xylose and RT. Contrast settings were the same as for TGP in figure 1C, additionally high contrast was used to show low levels of fluorescence.

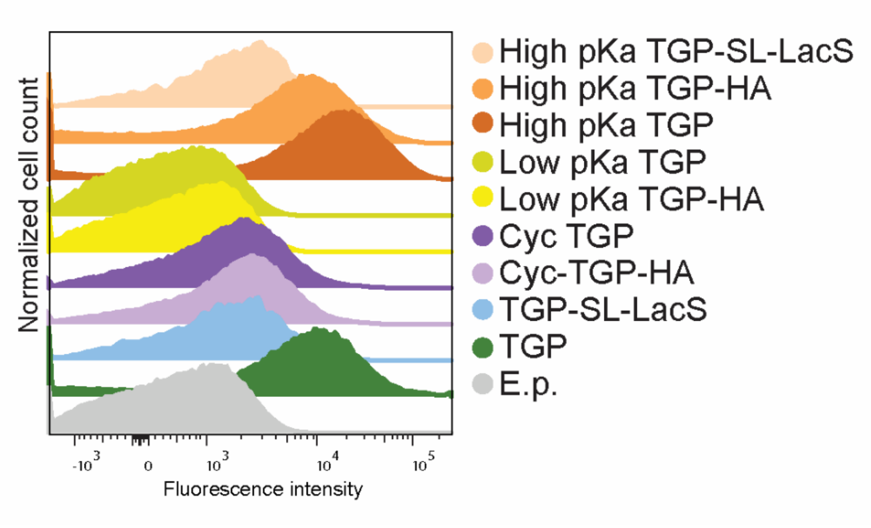

Figure S2. TGP variants with mutations introduced to modify pKa and sulfur bridges. Fluorescence intensity in flow cytometry at RT and 0.2% xylose. Number of events per sample: 100.000.

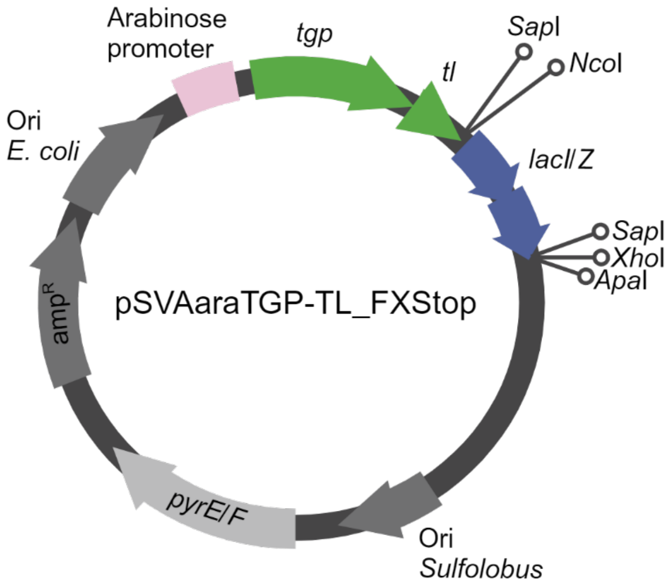

Figure S3. Schematic design of the plasmid pSVAaraTGP-TL_FXStop (pSVA6514) containing an arabinose promoter, *tgp,* the thermolinker (*tl*) and *lacI/Z* flanked by restriction sites *Nco*I and *Xho*I to inserte a gene of interest in frame with TGP-TL (thermolinker), followed by a stop codon. Alternatively, *Sap*I can be used to clone via FX cloning. Amp$^{R}$: ampicillin resistance cassette for selection in *E. coli*, *pyrE/F*: cassette for selection in Sulfolobus (uracil auxotrophy), Ori: origin of replication.

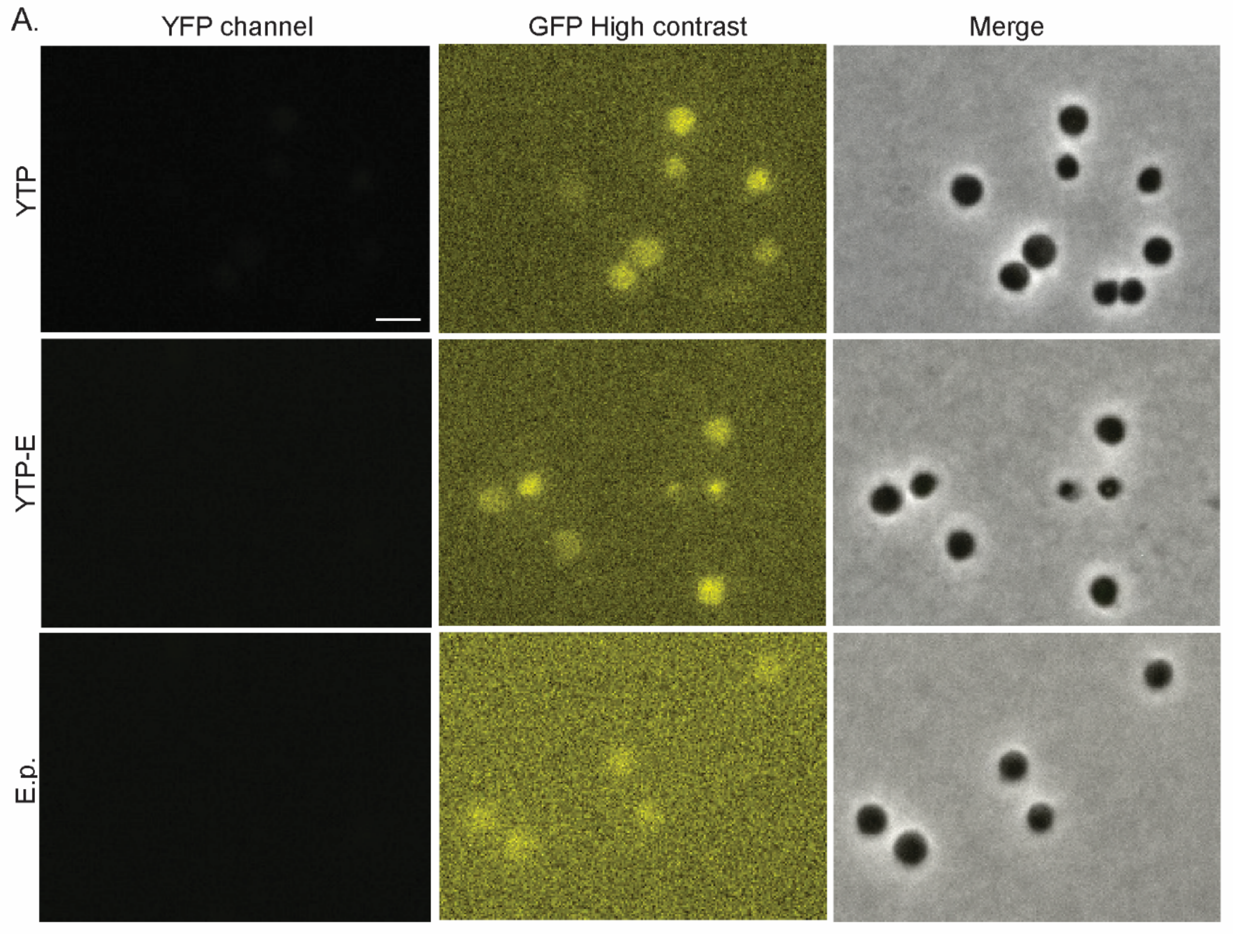

Figure S4. Microscopy images of yellow FPs. Fluorescent microscopy, phase contrast and merge with phase contrast images of cells expressing YTP and YTP_E. Contrast settings were the same as for mfYFP in figure 4, additionally high contrast was used to show low levels of fluorescence. E.p.: Empty plasmid

**Table S1.** Plasmids used in this work

| pSVA | pSVA backbone | Description | Reference |
|---|---|---|---|
| araFx_Stop | | Backbone for expression plasmids | (van der Kolk et al., 2020) |
| araFX_HA | | Backbone for expression plasmids | (van der Kolk et al., 2020) |
| araFX_NtHA | | Backbone for expression plasmids | (van der Kolk et al., 2020) |
| **Green fluorescent proteins** | | | |
| 12825 | pSVAaraFx_Stop | TGP | This work |
| 2584 | pSVAaraFX_NtHA | HA-TGP | This work |
| 2585 | pSVAaraFX_HA | TGP-HA | This work |
| 5597 | pSVA2585 | LacS-SL-TGP-HA | This work |
| 13120 | pSVAaraFX_HA | sfGFP-HA | This work |
| 13127 | pSVAaraFX_HA | eCGP123-HA | This work |
| 13157 | pSVA12825 | TGP-SL-LacS | This work |
| 13186 | pSVA2585 | TGP-SL-LacS-HA | This work |
| 6514 | pSVAaraFX-Stop | pSVAaraTGP-TL_FXStop | This work |
| **Linkers** | | | |
| 13158 | pSVAaraFx_Stop | TGP-TL-LacS | This work |
| 13159 | pSVAaraFx_HA | TGP-TL-LacS-HA | This work |
| 13184 | pSVA2585 | LacS-TL-TGP | This work |
| 13143 | pSVAaraFx_HA | pSVAara_FL_TGP_HA | This work |
| 13144 | pSVAaraFx_HA | pSVAara_RL _TGP_HA | This work |
| 13145 | pSVAaraFx_HA | pSVAara_SRL_TGP_HA_ | This work |
| 13146 | pSVA13144 | LacS-RL-TGP-HA | This work |
| 13148 | pSVAaraFx_Stop | LacS-RL-TGP | This work |
| 13151 | pSVA13145 | LacS-SRL-TGP-HA | This work |
| 13154 | pSVAaraFx_Stop | LacS-SRL-TGP | This work |
| 13155 | pSVA13143 | LacS-FL-TGP-HA | This work |
| 13156 | pSVAaraFx_Stop | LacS-FL-TGP | This work |
| **TGP variants** | | | |
| 13178 | pSVAaraFx_Stop | Cyc TGP (C103S C113S C171S) (codon adapted gene ordered in GenScript) | This work |
| 13179 | pSVAaraFx_Stop | Low pKA TGP (K5E K9E K21E K137E R186E K202E) (codon adapted gene ordered in GenScript) | This work |
| 13180 | pSVAaraFx_Stop | High pKA TGP (E26R E45R E73R E89R E109R E117R D187R) (codon adapted gene ordered in GenScript) | This work |
| 13181 | pSVAaraFX_HA | Cyc TGP-HA | This work |
| 13182 | pSVAaraFX_HA | Low pKa TGP-HA | This work |
| 13183 | pSVAaraFX_HA | High pKA TGP -HA | This work |

| | | | |
|---|---|---|---|
| 13187 | pSVA13183 | High pKa TGP-SL-LacS-HA | This work |
| **Cell localization** | | | |
| 6500 | pSVAaraFX_Stop | Expression TGP-TL-cdvB (saci_1373) | This work |
| 6552 | pSVAaraFx_Stop | native promoter TGP-TL-PCNA1 (saci_0826) | This work |
| **Yellow fluorescent proteins** | | | |
| 13126 | pSVAaraFX_HA | YFP-HA | This work |
| 6506 | pSVAaraFX_Stop | hfYFP (codon adapted gene ordered in GenScript) | This work |
| 6545 | pSVAaraFX_Stop | mfYFP | This work |
| 6593 | pSVAaraFX_HA | mfYFP-HA | This work |
| 6594 | pSVAaraFX_NtHA | HA-mfYFP | This work |
| 6586 | pSVA12825 | YTP | This work |
| 6576 | pSVA6586 | YTP-E | This work |

**Table S2.** Primers used in this study

| Primer No. | Sequence (3´->5´) | Use |
|---|---|---|
| 9355 | GCGCCTCGAGTCCACCACCACTACCACC | Rv *XhoI tgp* |
| 9356 | GCGCCCATGGCGGCAAGTGTAATAAAGCCTGAGATG | Fw *NcoI tgp* |
| 8662 | GACTCCATGGACTCATTTCCAAATAGCTTTAGG | Fw *lacS* |
| 8663 | GACTGGATCCGTGCCTTAATGGCTTTACTGGAGGTACGCTAT | Rv *lacS* |
| 11657 | ATATATGCTCTTCTAGTTCTAAGGGTGAGGAGTTATTCACTGGA | Fw *Sap*I *sfGFP* |
| 11658 | TATATAGCTCTTCATGCCCTTATTTATATAACTCATCCATTCCA | Rv *Sap*I *sfGFP* |
| 11680 | ACGTCCATGGATGAGTGTAATAAAGCCTGAAATGA | Fw *Nco*I *eCGP123* |
| 11681 | GCATCTCGAGTTTAGCTTGACTTGGTAACATA | Rv *Xho*I *eCGP123* |
| 11682 | ACGTCCATGGATGTCTAAGGGTGAAGAGTTAT | Fw *yfp* |
| 11683 | GCATCTCGAGTTTATATAACTCATCCATTCCA | Rv *yfp* |
| 12026 | GACTCCATGGACTCATTTCCAAATAGCTTTAGGTTT | Fw *Nco*I *lacS* |
| 12027 | GACTCCATgGGAAATTCACCTGATCCAGCAGCTGAACCAGCTGATCC GTGCCTTAATGGCTTTACTG | Rv *XhoI* lacS |
| 12045 | ATGGCGGCAAGTGTAATAAAGCCT | Fw plasmid FX TGP |
| 12046 | CTCGAGTGCTGAAGAGCCGGATCC | Rv plasmid FX TGP |
| 12047 | GGATCCGGCTCTTCAGCACTCGAG | Fw fragment pUC linkers |
| 12048 | AGGCTTTATTACACTTGCCGCCAT | Rv fragment pUC linkers |
| 12058 | GCAGCAGCTTCCTTTGCAGCTGCCTCTTTAGCTGCAGCTTCCTCGAGTGC TGAAGAGCCGGAT | Fw rigid linker T4PKN |
| 12059 | AAAAGAGGCAGCTGCTAAGGAAGCAGCAGCTAAAATGGCGGCAAGTGTAA TAAAGCCTGA | Rv rigid linker T4PKN |
| 12062 | GCATCTCGAGGGATCAGCTGGTTCAGCTGCTGGATCAGGTGAATTTATGGA CTCAT TTCCAAATAGCTTTAGG | Rv *XhoI* lacS |
| 12071 | CATGCATGGGCCCTTACTCAAGCGATCCACCACCA | Rv *lacS* thermolinker |
| 12073 | GCATCTCGAGATGGACTCATTTCCAAATAGCTTTAGG | Fw *XhoI* LacS |
| 12074 | GGATCAGCTGGTTCAGCTGCTGGATCAGGTGAATTTATGGACTCATTTCCAA ATAGCTTTAGG | Fw LacS thermolinker |
| 12075 | AAATTCACCTGATCCAGCAGCTGAACCAGCTGATCCTCCACCACCACTAC CACCACCAGA | Rv TGP thermolinker |
| 12088 | CATGCCATGGGAGCACATGCTTCTGTAA | Fw *Nco*I *tgp* |
| 12092 | ATGGCGGCAAGTGTAATAAAGCCT | Fw to put pSVA13184 in frame |
| 12093 | AAATTCACCTGATCCAGCAGCTGAA | Rv to put pSVA13184 in frame |
| 13216 | ATGCCTCGAGTGCTCCTCCGCCACTTCCTCCACCA | Rv TGP variants |
| 13247 | GGATCAGCTGGTTCAGCTGCTGGATCAGGTGAATTTATGTTTGATAAGTTAT CGA TAATT | Fw *cdvB* thermolinker |
| 13248 | TGCATCTCGAGACCCTCAAGAACAATTAGACCCTT | Rv *Xho*I *cdvB* |
| 13281 | ATGCCCATGGTATCAAAGGGTGAAGAAC | Fw *Nco*I hfYFP/mfYFp |
| 13282 | TGCATCTCGAGTTTATATAATTCATTCATATCATGAGTT | Rv *Xho*I hfYFP |
| 13964 | CATAATGTTTATATAACAGCAG | Fw point mutation S147P in hfYFP |
| 13965 | AGGATTGAAATTATATTCTAACTTATG | Rv point mutation S147P in hfYFP |
| 13966 | CCTGATAATCATTATCTAAGTTATCAAT | Fw point mutation L195M in hfYFP |
| 13967 | CATTAGTACTGGTCCATCACCTATT | Rv point mutation L195M in hfYFP |
| 13968 | AAACTATCTAAGGATCCTAATGAGAAA | Fw point mutation V206K in hfYFP |

| | | |
|---|---|---|
| 13969 | TGATTGATAACTTAGATAATG | Rv point mutation V206K in hfYFP |
| 13981 | GGATCAGCTGGTTCAGCTGCTGGATCAGGTGAATTTATGATAAAAAATA AAGTTATG TCAGAT | Fw thermolinker *pcna1* |
| 13982 | TGCATCTCGAGAGATAGTTTGGGTGCAAGTAAATA | Rv *Xho*I *pcna1* |
| 13983 | ATCGCCGCGGTACCTTATATCTCTTCGATAGTAC | Fw *Sac*II Native promoter PCNA2 |
| 13984 | GACTCCATGGGAGGTTTCCCCGTATTATTATT | Rv *Nco*I Native promoter PCNA1 |
| 14083 | TATGAAGTTGATCATAGAATAG | Fw point mutation H193Y in TGP |
| 14084 | AGCATCTGGTAATCTAACATC | Rv point mutation H193Y in TGP |
